# Community turnover by composition and climatic affinity across scales in an alpine system

**DOI:** 10.1101/659169

**Authors:** Brian V. Smithers, Meagan F. Oldfather, Michael J. Koontz, Jim Bishop, Catie Bishop, Jan Nachlinger, Seema N. Sheth

## Abstract

**Premise of the study:** Examining community turnover across climate gradients at multiple scales is vital to understanding biogeographic response to climate change. This approach is especially important for alpine plants in which the relative roles of topographic complexity and non-climatic or stochastic factors vary across spatial scales.

**Methods:** We examined the structure of alpine plant communities across elevation gradients in the White Mountains, California. Using community climatic niche means (CCNMs) and measures of community dissimilarity, we explored the relationship between community composition and elevation gradients at three scales: the mountain range, individual peaks, and within elevation contours.

**Key Results:** At the mountain range scale, community turnover and CCNMs showed strongly significant relationships with elevation, with an increase in the abundance of cooler and wetter-adapted species at higher elevations. At the scale of a single peaks, we found weak and inconsistent relationships between CCNMs and elevation, but variation in community composition explained by elevation increased. Within the elevation contours, the range of CCNMs was weakly positively correlated with turnover in species identity, likely driven by microclimate and other site-specific factors.

**Conclusions:** Our results suggest that there is strong environmental sorting of alpine plant communities at broad scales, but microclimatic and site-specific, non-climatic factors together shape community turnover at finer scales. In the context of climate change, our results imply that community-climate relationships are scale-dependent, and predictions of local alpine plant range shifts are limited by a lack of topoclimatic and habitat information.

## INTRODUCTION

Mountain environments contain important ecosystems and are heralds of species distribution shifts in response to climate change (Körner, 2003). Pronounced warming at higher elevations exacerbates this potential for mountains to exhibit early signs of biotic responses to climate change (Pepin et al., 2015). Species have responded by generally moving upslope in response to warming (Parmesan and Yohe, 2003; Lenoir et al., 2008; Pauli et al., 2012; Millar et al., 2015; Smithers et al., 2018; Malanson et al., 2019), with this upslope advance having accelerated in recent years (Steinbauer et al., 2018). However, it is increasingly recognized that climate change expectations of wholesale upward or poleward shifts of species distributions are overly simplistic (Lenoir and Svenning, 2015), especially in topographically complex landscapes (Rapacciuolo et al., 2014). Further, alpine species, often cool-adapted and long-lived, may be both directly sensitive to temperature change due to their physiology and indirectly sensitive due to competitive exclusion by species recruiting from lower elevations (Alexander et al., 2015; Rumpf et al., 2018). Predicting how these direct and indirect impacts of climate change will together shape future communities requires a better understanding of how complex mountain landscapes shape community assembly.

A long-standing debate in community ecology is whether community assembly is repeatable and predictable. If environmental filtering shapes community assembly, then there should be repeated, deterministic patterns of community composition across similar climatic and geologic conditions (Clements, 1916, 1936). On the other hand, less predictable, or stochastic, processes in the form of chance colonization, random extinction, ecological drift, and dispersal limitation could be more important in explaining patterns of community assembly (Gleason, 1927; Slatkin, 1974). While this debate has often been waged as either deterministic or stochastic forces being the driver of community assembly, it is more likely that communities are structured by both processes (Chase and Myers, 2011). In the context of climate change, this debate can be further distilled to a debate about whether climatic or non-climatic factors structure community assembly. The relative importance of deterministic climatic factors versus deterministic non-climatic or stochastic factors (hereafter referred to as non-climatic factors) can change depending on the system or scale at which we examine it. In alpine systems, the relative importance of these factors in shaping plant communities is particularly unclear, and this knowledge gap limits our ability to predict the rate and magnitude of plant community change in alpine systems in response to climate warming.

A first step to understanding the relative importance of these factors in shaping plant communities is to examine how communities are currently distributed across multiple scales of climatic complexity. In mountain systems, scales for describing plant community response to climate change can range from a single elevation at one location to an entire mountain range. Elevation is a useful geographic feature when studying deterministic and stochastic effects on community assembly because at different scales of elevation range, the relative importance of each process may change. Across an entire mountain range, the physical isolation of mountaintops may amplify the stochastic processes of dispersal and colonization as low-lying areas between suitable high-elevation habitat can create geographic barriers to species movement (Dirnböck et al., 2011). However at the scale of a single peak where dispersal is likely not limiting, the deterministic process of environmental filtering may play a larger role with high community turnover across steep climate gradients driven by elevation (Körner, 2003, 2007). Lastly, small-scale topographic complexity in the alpine zone can also drive variation in community composition (Van de Ven et al., 2007; Scherrer and Körner, 2011; Lenoir et al., 2013; Winkler et al., 2016). At this microclimatic scale, community variation within an elevation contour may be high where topographic complexity is high; the variety of potential habitats may deterministically facilitate close proximity of species and communities with a variety of climatic niches. However, within an elevation contour, other factors such as the geology, habitat stability (e.g., scree moves more than bedrock) or species interactions may also play a large role in community variation (Graae et al., 2018).

Comparing different types of community characteristics allows us to disentangle how climatic and non-climatic processes are shaping the community across scales. Community climatic niche affinities are metrics of the community (similar to any community-weighted mean traits) derived from climatic conditions found across the component species’ regional distributions (Landolt et al., 2010; Scherrer and Körner, 2011; Lenoir et al., 2013). At broad scales, plant species ranges are largely determined by a suite of environmental factors, with many of those factors climatically based. The suite of climatic factors that a species can tolerate can be referred to as its climatic niche affinity, and climatic tolerances of the community can be described using the combined climatic niche affinities of its component species.

When considering community climatic niche affinities, environmental filtering processes may drive species turnover. Conversely, stochastic processes such as the timing of species arrival into the community may drive community similarity metrics based on species composition alone (Fukami et al., 2005). Similar patterns of climatic niche affinities and community composition across elevation for a single scale would indicate that deterministic processes are driving much of the community turnover.

For a counter example, if the range of climatic niche affinities encompassed within elevation contours is not correlated with turnover in the community composition (ie., where species turnover is high, the range of climatic niche affinity values among species is low), this points to non-climatic factors shaping the community variation at this scale. These types of comparisons could be scaled up to look at different elevation bands on a single peak, among peaks in a mountain range, or even among mountain ranges.

Management of large areas of public lands for resilience with a changing climate challenges us to understand how communities will shift both locally and regionally (Pecl et al., 2017). Examining relationships between plant communities and climate gives key insights into the potential for species and communities to respond to climate change by tracking their climatic niche across different scales and may inform whether moving upslope in elevation is a predictable method for species to remain in their climatic niche in response to climate change. Evaluating changes in biodiversity and species distributional shifts requires long-term monitoring across multiple scales (Gottfried et al., 2012). The Global Observation Research Initiative in Alpine Environments (GLORIA) is an international collaboration assessing global distributional shifts of alpine plant species in response to changing climate (http://www.gloria.ac.at/). This international program was founded to provide cost-effective, universal monitoring protocols and a unifying network to investigate the rate and magnitude of the changes in alpine communities through time and spatial patterning at local, regional, and global scales (Grabherr et al., 2010). This network has led to key findings about the vulnerability of montane systems to changing climate, including upward distributional shifts of many alpine species and loss of cool-adapted species in mountains (Pauli et al., 2007, 2012; Gottfried et al., 2012; Lamprecht et al., 2018; Rumpf et al., 2018; Steinbauer et al., 2018). The North American chapter was founded in 2004 (Millar and Fagre, 2007) with installations in California and Montana, and now has installations in many mountain regions of North America. As part of this effort, GLORIA Great Basin (www.gloriagreatbasin.org) established a series of downslope transects on five peaks in the White Mountains, California (Fig. 1) extending from alpine mountaintops into the subalpine zone to monitor community changes in the local species pool at the individual mountain and at the mountain range scale.

**Figure 1.**
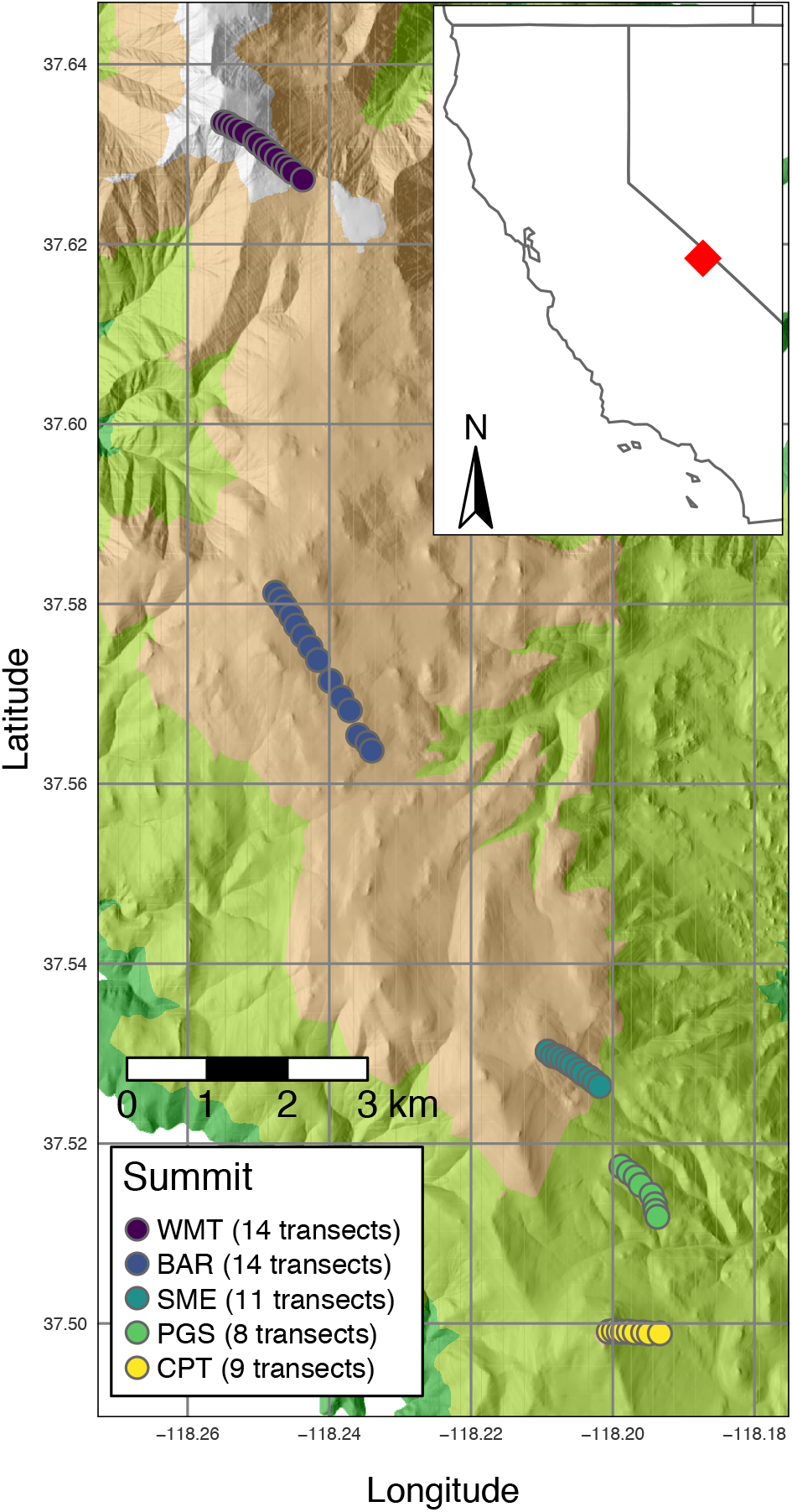
Geographic extent and terrain of the study area in the White Mountains, east of the Sierra Nevada mountain range in California (red diamond in inset). Points mark the centers for each transect, with 50m long and 1m wide belt transects extending on either side of these center points following the elevation contour. Terrain is rendered as a hillshade using the ⅓ arc second (~10m) digital elevation model from the USGS National Elevation Dataset with a sun angle of 40 degrees and a sun direction of 270 degrees. WMT = White Mountain Peak, BAR = Mount Barcroft, SME = Sheep Mountain East, PGS = Patriarch Grove South, and CPT = Campito Mountain. All transects are centered along the southeast aspect of each peak, except Campito Mountain (CPT), which is centered on the east aspect.

In this study, we evaluate the role of climatic versus non-climatic processes in explaining plant community turnover across elevation gradients at three scales in the White Mountains, California, USA: across an entire mountain range, within each of five peaks spread across the mountain range, and within elevation contours of each peak (Fig. 1). We use metrics of community climatic niche means and community composition to examine relationships among species turnover, community niche affinities, and elevation across scales. Consistent with environmental filtering, community climatic niche affinities may be highly dependent on the elevation gradient, with the presence and abundance of cooler and wetter-adapted species (dependent on their climatic niche) increasing with elevation, especially with climate metrics that vary the most predictably across the elevation gradient, such as temperature and precipitation. For this system specifically, previous work has found that community climate affinities are representative of local climatic niche means derived from microclimate field data (Oldfather and Ackerly 2019). However, given that the relative importance of deterministic and stochastic processes may vary with spatial scale and the level of community organization under consideration (Fukami et al., 2005), the relationships between elevation and climatic niche affinities may be strongest at broader spatial scales, while the relationship between elevation and community composition may be stronger at smaller scales (i.e., within a single peak). Specifically, we propose the following hypotheses (see Fig. 2 for a conceptual diagram).

1. At broad spatial scales (mountain range), non-climatic or stochastic processes will drive more variation in species composition and turnover than climatic or deterministic processes. Based on this, we predict that elevation as a proxy for deterministic changes explains more variation in species composition and turnover within each peak than across the entire range.
2. At broad spatial scales, climatic filters will be more important in explaining plant community variation in climatic niche space than non-climatic or stochastic differences. We predict that elevation will be better at predicting community climatic niche affinities at large spatial scales (mountain range) than at smaller scales (individual peak), where microclimate within an elevation may obscure this relationship
3. At the smallest scales (within elevation contour on a single peak), microclimate, habitat differences, and other unmeasured factors facilitate niche partitioning, thereby driving patterns of community assembly. We predict a positive relationship between the range of community climatic niche affinities and species turnover within elevation contours.

**Figure 2.**
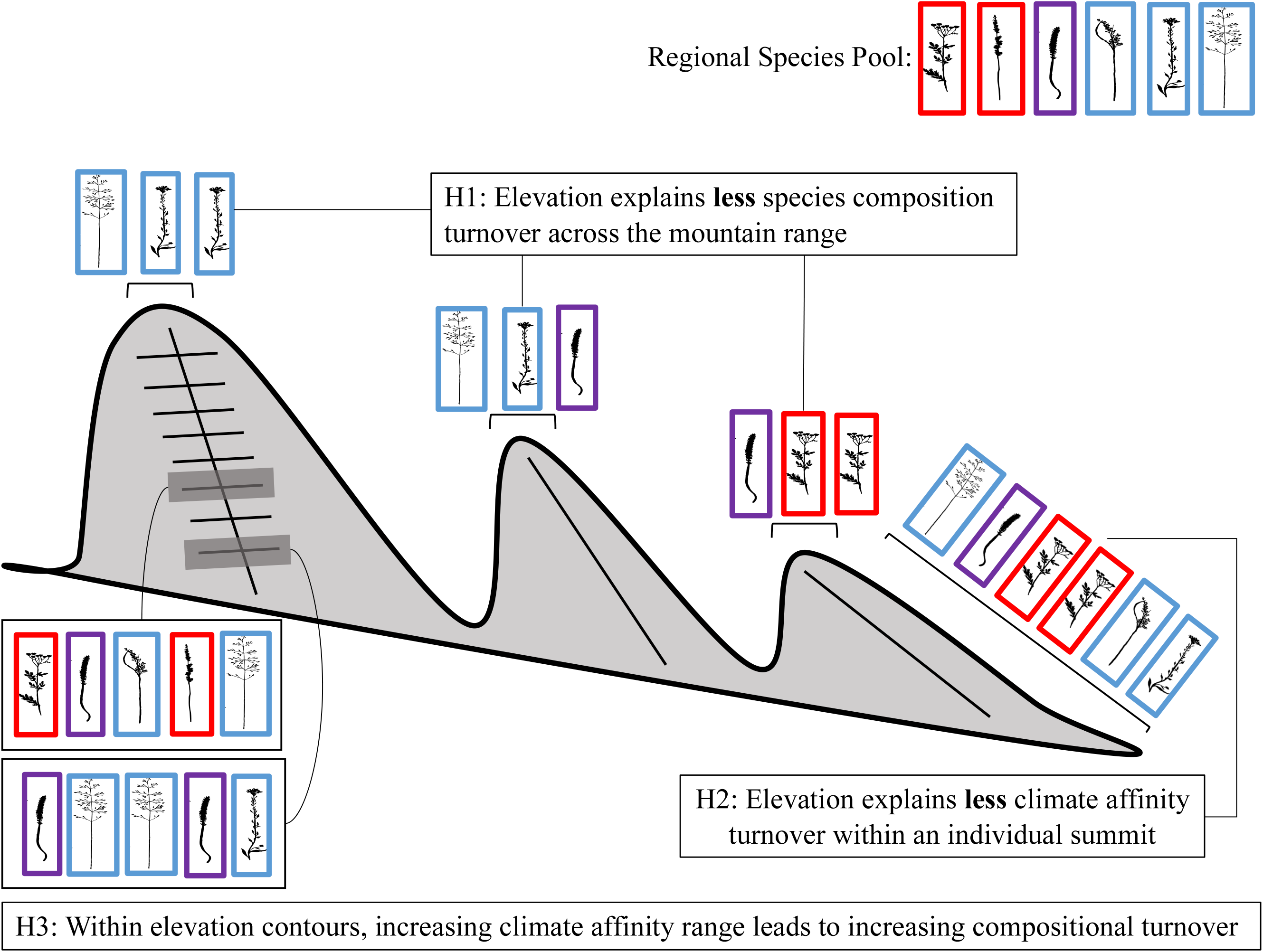
Conceptual figure illustrating the three presented hypotheses and the three scales of analysis: mountain range, summit, and within elevation contours. Species identity is represented by differing plant morphology and climatic niche affinity is represented by color. The first and second hypotheses (H1, H2) require comparisons of the general patterns of community turnover across the elevation gradient (for species identity and climatic niche affinity) between the mountain range and summit scales. Across the mountain range, we hypothesize predictive changes in the climatic niche affinity, but less so in species identity, across the elevation gradient. Within an individual summit, we hypothesize predictive changes in species identity, but less so in the climatic niche affinity of those species, across the elevation gradient. The third hypothesis (H3) focuses on comparing communities within elevation contours (single scale) and we show the hypothesized positive correlation between the turnover in species identity and range of climatic niche affinity.

## METHODS

### Study Area

The White Mountains are a cold and dry mountain range along the California-Nevada border at the western margin of the Great Basin in the eastern rain-shadow of the Sierra Nevada (Powell and Klieforth, 1991). At the White Mountains Research Center’s Barcroft Station (3800 m), mean annual precipitation is approximately 455 mm and mean annual temperature is −1.7 °C (Western Regional Climate Center). Most of the precipitation in the range comes as winter snow, although monsoonal rain events also occur during the summer. The alpine zone ranges in elevation from treeline at roughly 3500 m to the top of White Mountain Peak at 4300 m. The alpine flora in the White Mountains is made up of obligate alpine species, lower montane species, and cold-desert species associated with the Sierra Nevada, Great Basin, and Rocky Mountains (Rundel et al., 2008).

### Field Surveys

We sampled alpine community composition along elevation gradients using 100-meter belt transects on five peaks in the White Mountains: White Mountain Peak (WMT), Mount Barcroft (BAR), Sheep Mountain East (SME), Patriarch Grove South (PGS), and Campito Mountain (CPT) (Bishop, 2011). On each peak, the first transect was established on the southeast aspect (east aspect for Campito Peak), 35 vertical meters below the summit. Two, 50m x 1m belt transects were extended from this central point following the elevation contour to create a 100-meter x 1-meter belt transect at one elevation. Additional 100m x 1m transects were established on each peak in intervals of 25 vertical meters below the previous transect on the same aspect (Fig. 1). While transects were placed at this fixed 25 m interval, peaks had differing numbers of transects based on the total distance between upper treeline and the top of the peak. Transects were established across 3 consecutive years from 2011-2013 (Table 1). Sampling was conducted in mid-July to best correspond with flowering timing of most alpine plant species in the White Mountains. Each transect was divided into ten continuous, 10m x 1m segments. In each segment, we recorded the presence of every species. In addition, we assessed cover of plant species and substrate (rock, scree, litter, bare ground) along the entire transect using a point-intercept method with 40 points per 10m x 1m segment. A pair of points was located at every half-meter along the transect, with the points in each pair 0.5 m apart (0.25 m from the center of the belt transect). The substrate or plant species was recorded as a “hit” if a point intersected with it.

**Table 1:** Sampled peaks and associated data in the White Mountains, CA. Total richness is the number of unique species found on each peak.

| <u>Peak</u> | <u>Sample</u><br><u>Year</u> | <u>Elevation (m)</u> | <u>Soil Type</u> | <u>Aspect</u> | <u># Transects</u> | <u>Richness</u> |
| --- | --- | --- | --- | --- | --- | --- |
| White Mountain Peak (WMT) | 2012 | 3971-4296 | Igneous | SE | 14 | 50 |
| Mount Barcroft (BAR) | 2011 | 3625-3950 | Igneous | SE | 14 | 69 |
| Sheep Mountain (SME) | 2011 | 3475-3725 | Dolomitic | SE | 11 | 57 |
| Campito Mountain (CPT) | 2013 | 3325-3500 | Quartzitic | E | 8 | 51 |
| Patriarch Grove South (PGS) | 2012 | 3299-3478 | Dolomitic | SE | 8 | 51 |

### Community Metrics

To investigate how communities varied across spatial gradients, we used community-weighted climatic niche means (CCNM) and community composition. A CCNM is the abundance-weighted mean of each component species’ average species niche space across its range (De Frenne et al., 2013). For community composition we used the Bray-Curtis dissimilarity metric of *β* diversity, which incorporates both species presence and relative abundance (Anderson et al., 2011). Our measure of abundance used for both metrics was the number of segments in which it was found for each transect. We consider how these two community metrics change across three nested scales: the elevation gradient of the entire mountain range, the elevation gradient of a single peak, and within a single elevation contour of a peak.

A CCNM is a property of a single community, or in our specific case, a transect for mountain range- and peak-scale analyses and a 10m segment within transects for the elevation contour-scale analysis. We focused on the following climatic measurements as they are likely major structuring factors for alpine plant communities: climatic water deficit, mean annual precipitation, July maximum temperature, and January minimum temperature. Climatic water deficit is an integrative measure of how much energy availability (temperature) exceeds water supply (precipitation), with higher values indicating hotter, drier conditions, and lower values indicating cooler, wetter conditions. July maximum and January minimum temperatures were calculated mean daily high and low temperatures of the month, respectively. Using a recently compiled database of California herbarium specimens derived from the Consortium of California Herbaria as well as additional natural history museums and herbaria (Baldwin et al., 2017), we extracted locations from all vouchered specimens in the database for each surveyed species. We used those database locations to extract 30-year (1961-1990) means for climatic water deficit, mean annual precipitation, maximum July temperature, and minimum January temperature from the TerraClimate gridded dataset (Abatzoglou et al., 2018), which is itself an amalgamation of the WorldClim dataset (Fick and Hijmans, 2017) and other data. After culling the herbaria records to one record per species per grid cell to reduce sampling bias, we extracted the four mean climate values for each species, representing the mean climatic niche of the species for each climate metric (Appendix S1; see Supplemental Data with this article). This 30-year period was chosen as the temporal range of most likely establishment for the majority of the generally long-lived individual plants sampled. For each transect on each of the five peaks, we calculated a weighted mean of the mean climatic niche value of each species present in that transect. The mean was weighted by the abundance of each species in that community, in our case the number of segments in which it was found when the community was defined as the whole transect and the number of point-intercept hits when the community was defined as the 10m x 1m segments within the transect. The abundance weighting allows more dominant species to have a larger effect on the community’s CCNM. This final calculation produces one CCNM value for each climate metric in each community, allowing for examination of how the overall climatic niche of the community changes across the elevation gradients or ranges within elevation contours.

To examine community patterns within the elevation contours we used two metrics to represent how species identity and climatic niches vary across the contour: *β* diversity and CCNM range. We calculated per-transect *β* diversity as the total species richness in each transect divided by the mean species richness across the 10 segments within each transect (Whittaker, 1960; Anderson et al., 2011). Therefore, the minimum possible value is a *β* diversity of 1, which would indicate that every species in the whole transect is represented within each segment of that transect. This metric gives a measure of the amount of species variation that can be found within a single elevational band. The range of CCNM within a single transect was determined by calculating the CCNM per segment, and then subtracting the minimum from the maximum of each transect for each climate variable. The CCNM range gives a measure of how variable the climatic niches of the communities are along the elevation contour; a low range indicates that elevation is predominantly driving any variation in the climatic niche of the communities, and a high range indicates that microclimate may be shaping the patterns of community climatic niches across the landscape as well.

### Statistical Analyses

In order to quantify the community turnover explained by the elevational gradient at the scale of the mountain range and all peaks separately, we performed permutation-based MANOVAs using the ‘adonis’ function (1000 permutations) in the vegan package (Oksanen et al., 2008). We built separate MANOVAs for the entire mountain range and for each of the peaks. For each of these models, the dependent variable is a Bray-Curtis dissimilarity matrix (with the matrix rows representing single transects) and the independent variable is elevation across either the entire mountain range or across each peak, respectively. To visualize these relationships, we performed an ordination based on non-metric dimensional scaling (NMDS) that included community data from each peak across the entire White Mountain range.

To examine how CCNMs changed across the entire mountain range, we constructed linear mixed effects models for each of the four CCNMs (climatic water deficit, mean annual precipitation, maximum July temperature, and minimum January temperature) as the response variable, elevation as the predictor variable, and with a random intercept effect of peak. We compared models with and without a quadratic term for elevation and used Akaike’s Information Criterion (AIC) to choose the model with the best fit. We used a Bonferroni-corrected significance threshold of P=0.0125 for each of the four models representing the different CCNM responses across all the peaks to account for multiple testing. For each of the four CCNMs, we also built linear models for each peak with CCNM as the response variable and elevation as the predictor variable. We used a Bonferroni-corrected significance threshold of P=0.01 for each of the five models representing, for each CCNM, the CCNM response for the five different peaks.

To examine community composition and species turnover at the scale of each elevation contour, we used linear models to determine whether the range in CCNM within an elevation contour (transect) significantly explained variation in *β* diversity. All analyses were performed using R version 3.6.0 (R Core Team, 2018). Data manipulation and visualization were performed using the R packages dplyr (Wickham et al., 2019), sf (Pebesma, 2018), and ggplot2 (Wickham, 2016).

## RESULTS

### Community Composition

We observed a total of 123 species in 70 genera and 25 families across the entire mountain range, with individual peak species richness ranging from 50-69 (Table 1). Across the mountain range, elevation explained 30% of the variation in community composition (*P* < 0.001, Fig. 3). When looking at each peak independently, there was an approximate 10-20% increase in the community variation explained by elevation (BAR 42% *P* < 0.001, CPT 37% *P* = 0.012, PGS 40% *P =* 0.002, SME 42% *P* < 0.001, WMT 52% *P* < 0.001). The NMDS ordination visualizes these relationships by reducing the community dimensionality into 2 dimensions. These ordinations show a clear range-wide relationship between the first axis (NMDS1) and elevation (95%, *P* < 0.001), as indicated by the arrow. Therefore, species such as *Erigeron vagus* are associated with higher elevation sites and *Erigeron tener* is associated with lower elevation sites. Within the summits, the community composition had stronger, but unique relationships with elevation; the ellipses that encompass the standard deviation of points associated with each summit line up well with the first axis and vary across the second axis (NMDS2).

**Figure 3.**
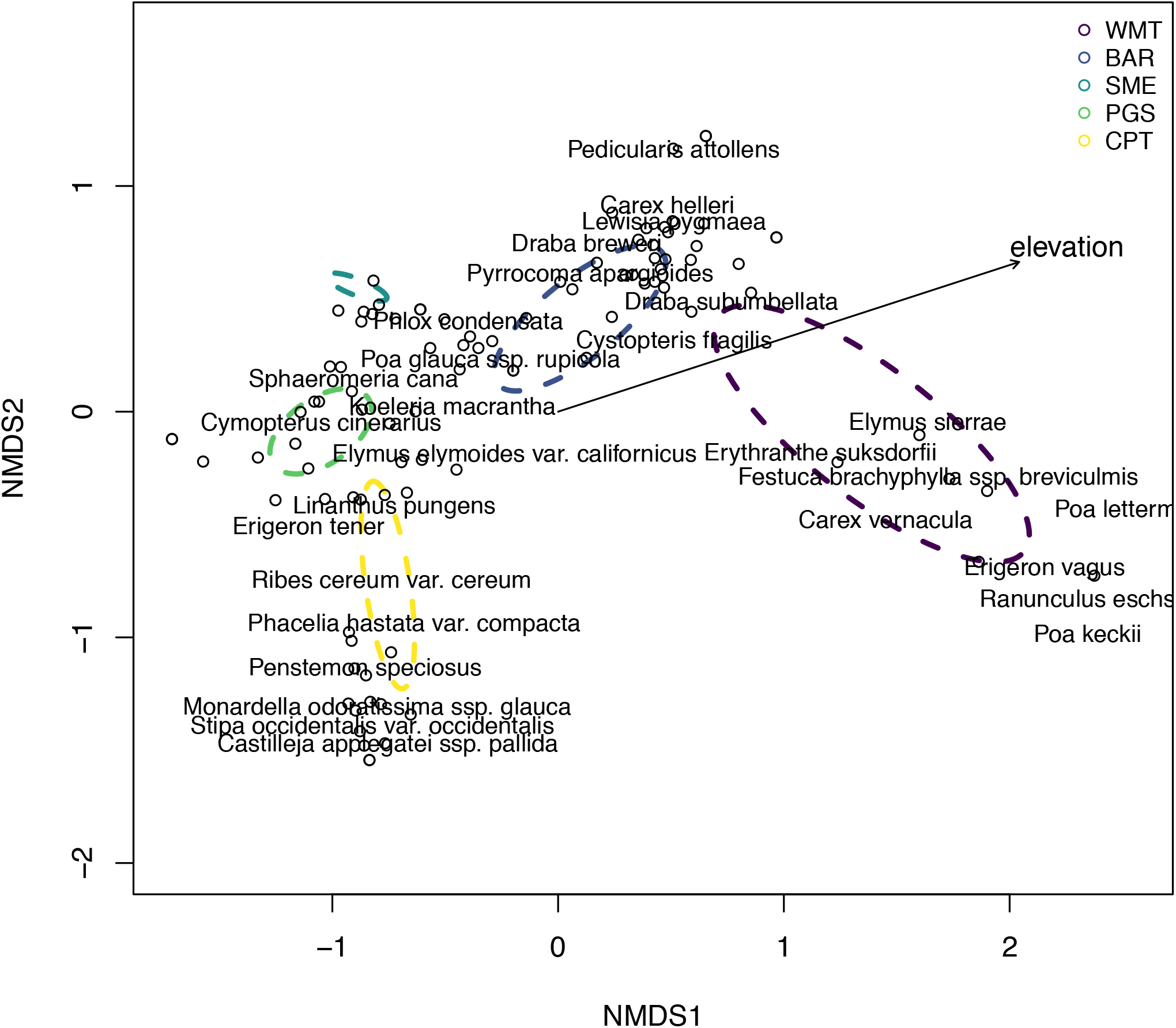
Non-metric dimensional scaling (NMDS) ordination of Bray-Curtis dissimilarity. The arrow indicates the directional influence of elevation. In the event of species name overlap, only the more common species is shown here, but all species were included in the analysis.

### Community-weighted Climatic Niche Mean

All CCNM metrics (climatic water deficit, mean annual precipitation, maximum July temperature, and minimum July temperature) showed highly significant linear relationships with elevation at the mountain range scale (*P* < 0.001 for all metrics, Fig. 4, Appendix S2). For each CCNM at the mountain range scale, the non-quadratic model performed best.

**Figure 4:**
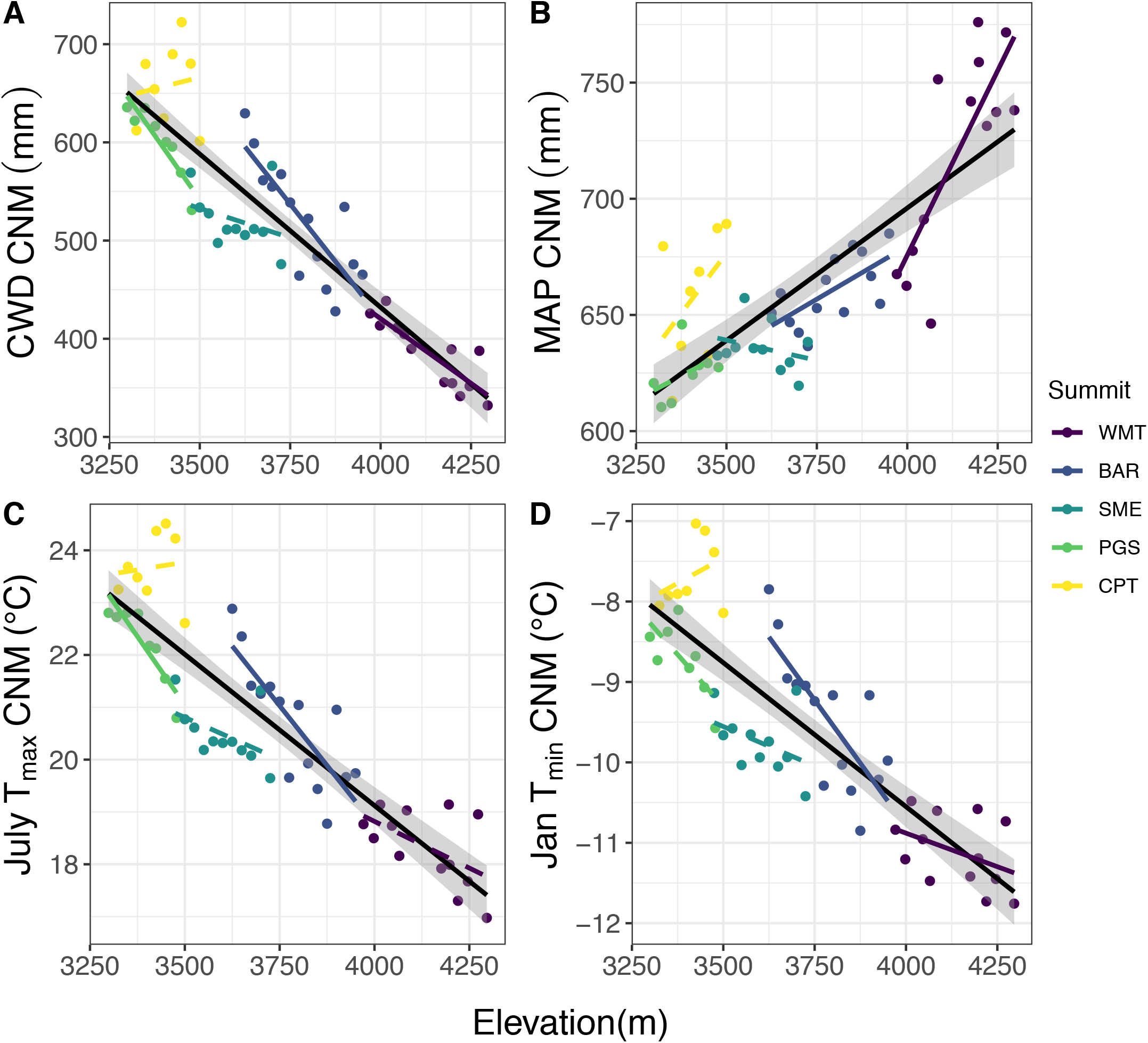
Community-weighted Climatic Niche Means for (A) Climatic Water Deficit (CWD), (B) Mean Annual Precipitation (MAP), (C) Average July maximum temperature (July T_max_), and (D) Average January minimum temperature (Jan T_min_) along the elevation gradient. The black solid lines represent the linear model fit for the entire range, all of which showed highly significant trends (*P* < 0.001). Shaded areas represent the standard error around the linear model. Each point is 1 transect and is color-coded by peak. Peaks with a significant (*P* < 0.05) relationship between CCNM and elevation are shown with a solid line, and those with a non-significant (*P* > 0.05) relationship are shown with a dashed line.

Climatic water deficit, maximum July temperature, and minimum January temperature CCNMs showed a negative relationship with elevation while mean annual precipitation showed a positive relationship with elevation. Collectively, these results indicate that the presence and abundance of species with cooler, wetter niches increased with elevation. At the individual mountain scale, the relationships between the CCNM and elevation were weak, inconsistent, or absent (Fig. 4, Appendix S2).

### Variation in community composition and CCNMs across elevation contours

The mean *β* diversity per transect across the elevation contours was 2.1 [range, 1.3-3.2]. A *β* diversity of 2 indicates that, on average across the segments of the transect, there are twice as many species represented in the whole transect than in each segment. The mean of the ranges of CCNM per transect across the elevation contours were climatic water deficit: 173 mm [0 – 465], mean annual precipitation: 15.3 mm [0 – 47.9], maximum July temperature: 1.61°C [0 – 3.88], and minimum January temperature: 2.01°C [0 – 5.75]. Interestingly, this range is slightly lower, but similar in magnitude as the range of CCNM values observed across the entire mountain range (Fig. 5).

**Figure 5.**
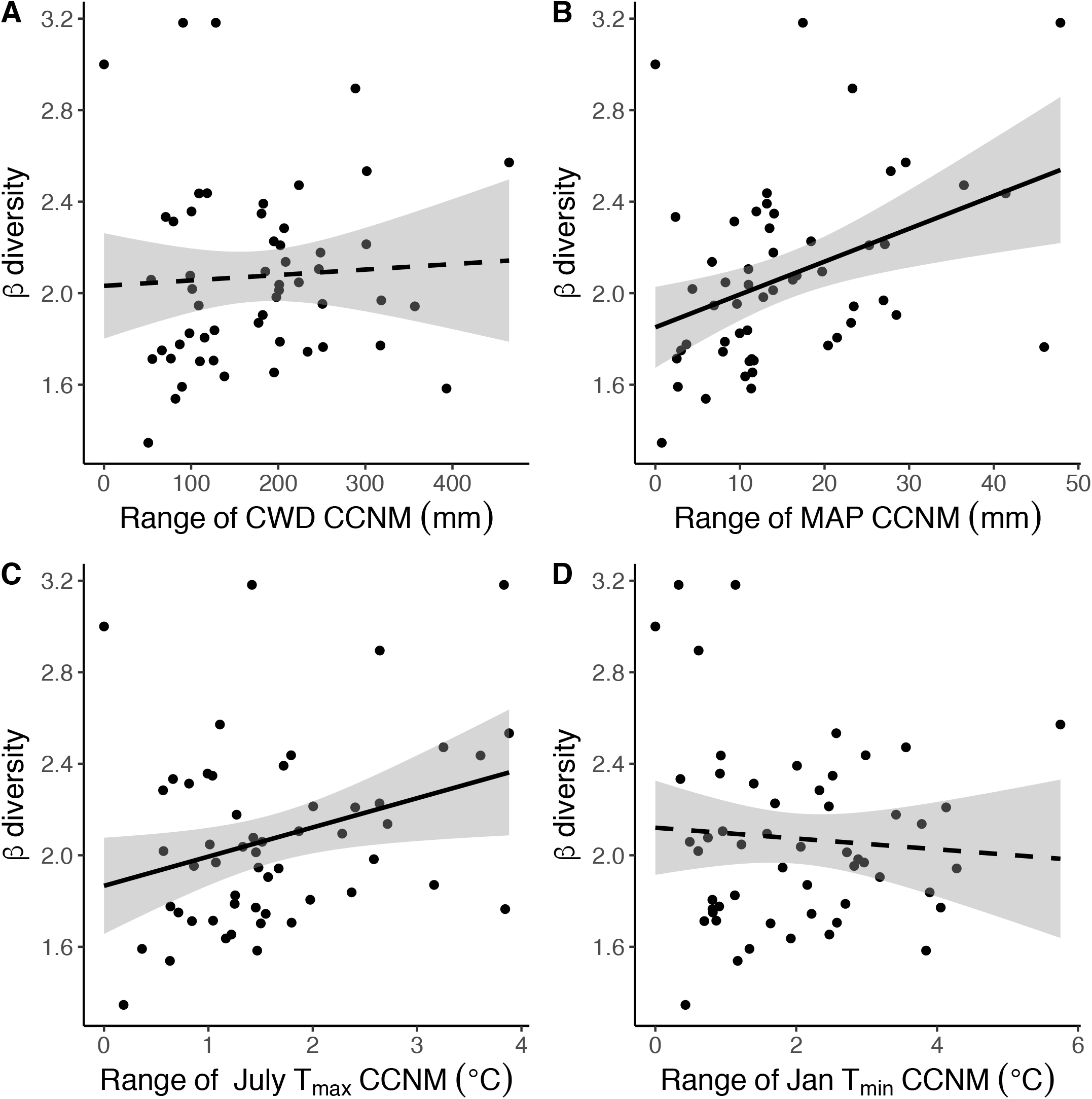
Relationship between community climatic niche mean (CCNM) range and per-transect *β* diversity. Climate metrics with a significant (*P* < 0.05) relationship between CCNM range and βdiversity per transect are shown with a solid line, and those with a non-significant (*P* > 0.05) relationship are shown with a dashed line.

For CCNMs based on mean annual precipitation and maximum July temperature, the CCNM range was significantly and positively correlated with the per-transect *β* diversity (mean annual precipitation: adjusted R^2^ = 0.138, *P* = 0.003; maximum July temperature: adjusted R^2^ = 0.073, *P* = 0.027; Fig. 5). For climatic water deficit and minimum January temperature, there was no significant relationship between CCNM range and per-transect *β* diversity (climatic water deficit: adjusted R^2^ = – 0.016, *P* = 0.683; minimum January temperature: adjusted R^2^ = −0.014, *P* = 0.590).

## DISCUSSION

Investigating the dynamics of community turnover across climate gradients is a fruitful avenue for understanding where and how species distributions may shift with a changing climate (Scherrer and Körner, 2011; Lenoir et al., 2013). Across the entire mountain range, both community metrics (climate-derived CCNM and composition-derived Bray-Curtis dissimilarity) exhibited strong relationships with elevation, but the relationships between CCNM and elevation at the individual peak scale was weak and inconsistent (Figs. 3, 4). In contrast, there was a stronger relationship between elevation and community composition at the scale of the individual peak relative to the entire mountain range (Fig. 3). Lastly, the range of community climatic niche means was positively, but weakly correlated with the per-transect *β* diversity (Fig. 5). Below, we discuss in more depth how the processes that shape alpine plant distributions and community assembly vary across scales and explore potential impacts for using these community metrics for predicting the effects of climate change on mountain communities.

### Community Composition and Turnover Across Scales

Elevation was a strong predictor of community dissimilarity and CCNM across the entire White Mountain range (Figs. 3, 4), suggesting that elevation structures both community composition and niche affinities across broad spatial scales. However, we found varying responses to elevation between the two community metrics at the scale of individual peaks. Consistent with our first hypothesis, more variation in community composition was explained by elevation at the individual peak scale relative to the entire range despite a lower elevation range within each individual summit relative to the mountain range. At the mountain range scale, non-climatic differences between peaks (e.g., biotic interactions or geology) or stochastic differences likely reduce the community variation explainable by elevation. Community dissimilarity has a strong, but unique relationship with elevation within each summit (Fig. 3), which muddles the broader-scale relationship between elevation and dissimilarity across summits.

At the scale of individual peaks, the elevation gradient may encompass variation in additional factors besides climate, such as available space, underlying changes in geology, biotic interactions, and microtopography. These non-climatic factors may influence patterns of species dissimilarity across the elevation gradients within each peak. Dispersal limitation may also weaken the relationship between elevation and community composition at the scale of the entire mountain range (Dirnböck et al., 2011). If stochastic processes such as dispersal shape community assembly, then community dissimilarity should not vary predictably across environmental gradients. But if deterministic processes such as environmental filtering shape community assembly, community dissimilarity will not vary predictably across spatial gradients (Chase and Myers, 2011). In our study, elevation encompasses both spatial and climatic gradients limiting our ability to assess the relative importance of stochastic vs. deterministic processes in shaping species (Fig. 1; Appendix S3). Nonetheless, if dispersal limitation is reduced within individual peaks relative to among peaks across the range, elevation may be more predictive of variation in community composition within peaks as species are filtered with respect to the climatic conditions across the gradient (Svenning and Sandel, 2013; Graae et al., 2018).

### Climatic Niche Means Across Scales

If climatic filtering is primarily responsible for shaping community assembly, then the presence and abundance of cooler, wetter-adapted species should increase with elevation, particularly at the scale of the entire mountain range. Indeed, we found that CCNMs based on all four climate metrics were significantly related to elevation at the mountain range scale (Fig. 4). Encouragingly, these relationships at this scale all also matched the predicted patterns of climatic niche means and elevation based on how the actual climate is predicted to vary across elevation gradients (Appendix S4). Species associated with cooler and wetter conditions were found more often and/or in higher abundance at higher elevations (Fig. 4). We also found support for our second hypothesis that elevation should more strongly predict climatic niche affinities at large spatial scales (mountain range) than at smaller scales (individual peak). Despite the strong, consistent relationship between CCNMs and elevation across the entire mountain range, dependent on the peak and the climate variable used to quantify CCNM, there were variable results for whether elevation was predictive of community niche affinities for individual peaks. Therefore, at finer scales, elevation is less consistently predictive of the climatic niche means of communities. This may be due to the reduced range of elevation change, and therefore climatic variation, within each summit, allowing non-climatic factors to play a larger role in community assembly. Within elevation contours, the range of community climatic niche affinities influenced variation in species turnover with the CCNMs based on July maximum temperatures and precipitation (Fig. 5), suggesting that for at least these climate metrics, microclimate may facilitate niche partitioning and species coexistence at the within-contour scale. Future studies should more formally examine the mechanisms of how microclimate drives patterns of community assembly across these three spatial scales.

One caveat of our methodology is that the peaks were surveyed over the course of three years. Interannual variation in weather could certainly have affected our ability to successfully identify all species present in the transects. While we attempted to survey during peak flowering of the majority of plant species in the White Mountains, it is also possible that different temperatures and precipitation amounts in the three years could have changed the species abundances or even shifted that peak phenological period outside of our sampling period. Since a given peak was surveyed in a single year, this potential issue could only affect the mountain range-scale analyses.

An important caveat of our use of CCNMs is that due to data availability, we quantified CCNMs with species’ occurrences from California distributions, not the species’ global distributions. For many species, California encompasses the range of climatic conditions that the species experience across their ranges. For other species, however, the occurrences in California are a subset of the species’ larger distribution, and hence their climatic niches. Niche estimates are likely particularly inaccurate for species whose western range limits are found in eastern California. Further, all CCNM calculations assumed no variation among populations across species ranges and thus do not account for the possibility that the populations surveyed in the White Mountains may have different climatic niches relative to those found in other parts of the species range. CCNMs also disregard influences of fine-scale climate on species’ distributions due to the coarse resolution of the climatic and herbarium specimen data (Bramer et al., 2018), potentially helping to explain the weaker and less consistent relationships found between elevation and CCNMs within each peak relative to across the entire mountain range.

### Drivers of Community Variation Within Elevation Contours

The maximum range in CCNMs observed within a single elevation contour was on par with the overall maximum range of CCNM values observed across the entire mountain range. This points to at least some role of microclimate in shaping species distributions and community assembly. Surprisingly, we found CCNM differences within elevation contours that were equivalent to those found across 100 m in elevation. While in general, species are expected to move upslope in response to warming, at the local scale there will likely be suitable microclimate within the same elevation. Species movement may be lateral or even downslope in order to track newly suitable microclimate. These findings are supported by research in alpine plant biology which has found that microclimate has a large influence on species’ distributions (Körner, 2003). However, the range of community climatic niche means was positively, but weakly, correlated with the *β* diversity within elevation contour (Fig. 5), indicating minor support for our third hypothesis.

Therefore, other non-climatic processes are also shaping the species turnover within the elevation contour. As dispersal limitation at this scale is unlikely, two major potential factors that may be instead be shaping community distributions are habitat type (e.g., presence of moveable scree) and species interactions. In addition to microclimate, substrate type (rock, scree, organic soil) likely heavily influence species distributions, as well as species’ responses to climate change (Kulonen et al., 2018). Species interactions directly, or in conjunction with habitat type and microclimate, can also influence species distributions at fine spatial scales (Alexander et al., 2015; Blonder et al., 2018; Graae et al., 2018). For example, lower elevation species may be more competitive on organic soils than on rock or scree whereas higher elevation obligate alpine plants potentially have more adaptive traits (e.g., contractile roots) for persisting on the dynamic scree habitat (Kulonen et al., 2018). As we build our temporal dataset, future studies will examine whether interactions between habitat segregation and elevation influence species and community responses to climate change over time.

### Conclusions/Implication for Climate Change Research

Our findings suggest that the effects of elevation on community niche affinities and community dissimilarity varies with spatial scale. At the scale of the entire mountain range, there is climatic filtering across the elevation gradient, with an increase in the presence and abundance of cool, wet-adapted species at higher elevations. Within individual peaks and elevation contours, however, the signal of climatic filtering breaks down. This points to differences in how climatic and non-climatic factors may shape species composition at different scales. At broad spatial scales, deterministic climate appears to be a main driver of plant community assembly since the large, consistent elevation gradient can mask stochastic factors such as chance colonization and extinction and non-climatic factors like geology. At smaller scales where dispersal is less limiting and microsite climatic variation is common even within the same elevation contour, synoptic climatic factors appear less important than non-climatic factors in structuring community. We did not directly measure those non-climatic factors, but they are likely to be related to habitat and substrate type.

Since climatic niche metrics are often used for quantifying community responses to climate change, it is important to understand their relative strengths in explaining community turnover across scales. The breakdown of correlations between CCNM and elevation at finer-scales, even though species turnover based on identity is still explained by elevation at this scale, suggests an important caveat for using climate-based community measures at these scales. Perhaps we would not find this pattern if our climate data used to calculate the species climatic niches were at finer resolutions, which is supported by the positive correlation per-transect βdiversity and range of CCNM within a transect. However, this correlation is weak indicating that higher resolutions may not be sufficient to understand species distributions at these micro-topographic scales. Within the context of climate change, predictions of local alpine plant range shifts will be limited by availability of topoclimatic, as well as habitat, information.

## Acknowledgements

The authors thank the UC Natural Reserve System’s White Mountain Research Center, USFS Pacific Southwest Research Station, Consortium for Integrated Climate Research in Western Mountains (CIRMOUNT), the Global Observation Research Initiative in Alpine Environments (GLORIA), and USFS Inyo National Forest for their support and collaboration. Our data collection teams consisted of many citizen scientists that include graduate, undergraduate, and high school students as well as members of NGOs such as GLORIA Great Basin, the California Native Plant Society, the Nevada Native Plant Society, and the Davis Botanical Society. We especially thank Connie Millar and Adelia Barber for their leadership and support of the early stages of this research. We would also like to thank two anonymous reviewers for their insightful comments. This work was supported by USDA National Institute of Food and Agriculture Hatch project 1016272.

## Author Contributions

Brian Smithers, Meagan Oldfather, and Michael Koontz participated in data collection, data processing, analysis, and in the writing of the manuscript.

Jim Bishop and Catie Bishop created the downslope transect protocol, participated in data collection, data processing, and in editing of the manuscript.

Jan Nachlinger participated in data collection, data processing, and in the editing of the manuscript. Seema Sheth participated in data processing, analysis and in the writing of the manuscript.

## Data Accessibility

Data and R code from the project are available on the GLORIA Great Basin website (https://www.gloriagreatbasin.org/documents-archives) as well as in a permanent Figshare repository (DOI: 10.6084/m9.figshare.8968793). Due to the long-term nature of plot monitoring, we have not collected voucher specimens from within the plots however when voucher specimens from outside the plot were collected, the specimens were housed at the White Mountain Research Center, Owens Valley Station.

## Supporting Information

Additional Supporting Information may be found online in the supporting information section at the end of the article.

Appendix S1: Climatic Niche Means (CNMs) for each species observed among all peaks.

Appendix S2: Model results for each of the climate metrics on each of the peaks and for all peaks combined.

Appendix S3: Average climate variables for each transect.

