## Appendix S1 for "Community turnover by composition and climatic affinity across scales in an alpine system"

##### Smithers et al.— bioRxiv 2019 – Appendix S1

##### Appendix S1: Climatic Niche Means (CNMs) for each species observed among all peaks.

| **Species** | **CWD (mm)** | **MAP (mm)** | **July T_max_ (°C)** | **Jan T_min_ (°C)** |
| --- | --- | --- | --- | --- |
| Agoseris monticola | 291.1 | 23.3 | 33.0 | -4.7 |
| Androsace septentrionalis | 188.4 | 13.4 | 35.2 | -3.8 |
| Anelsonia eurycarpa | 164.1 | 15.8 | 33.0 | -5.6 |
| Antennaria media | 234.6 | 26.2 | 32.7 | -4.4 |
| Antennaria rosea | 410.7 | 26.3 | 33.7 | -3.1 |
| Antennaria umbrinella | 272.5 | 15.8 | 34.7 | -4.0 |
| Artemisia dracunculus | 842.0 | 25.7 | 35.3 | 0.6 |
| Artemisia nova | 756.4 | 20.7 | 34.1 | -3.4 |
| Artemisia rothrockii | 407.7 | 13.7 | 35.9 | -1.5 |
| Astragalus calycosus var. calycosus | 489.3 | 14.9 | 32.4 | -5.9 |
| Astragalus kentrophyta var. danaus | 196.0 | 19.2 | 31.1 | -6.9 |
| Astragalus kentrophyta var. tegetarius | 268.7 | 15.9 | 31.8 | -6.3 |
| Astragalus lentiginosus var. semotus | 399.4 | 16.8 | 31.6 | -6.3 |
| Boechera depauperata | 239.2 | 16.7 | 33.5 | -5.2 |
| Boechera inyoensis | 379.4 | 15.2 | 33.2 | -5.8 |
| Boechera lyallii | 225.9 | 21.1 | 31.5 | -6.4 |
| Boechera pulchra | 993.5 | 21.0 | 35.8 | 0.3 |
| Calamagrostis purpurascens | 222.6 | 17.6 | 32.3 | -6.0 |
| Calyptridium roseum | 423.6 | 17.1 | 32.0 | -6.9 |
| Calyptridium umbellatum | 353.1 | 36.7 | 30.8 | -6.4 |
| Carex duriuscula | 160.0 | 13.0 | 33.5 | -6.6 |
| Carex haydeniana | 198.6 | 16.2 | 33.3 | -5.4 |
| Carex helleri | 203.2 | 17.0 | 32.9 | -5.5 |
| Carex incurviformis | 170.6 | 16.2 | 32.6 | -6.1 |
| Carex leporinella | 267.9 | 26.2 | 31.6 | -5.9 |
| Carex rossii | 427.1 | 41.2 | 31.3 | -4.8 |
| Carex subnigricans | 259.1 | 16.5 | 32.9 | -5.6 |
| Carex vallicola | 405.0 | 17.2 | 32.4 | -6.6 |
| Carex vernacula | 244.1 | 19.0 | 32.4 | -5.8 |
| Castilleja applegatei subsp. pallida | 303.7 | 20.9 | 31.4 | -6.4 |
| Castilleja linariifolia | 729.2 | 20.6 | 33.9 | -3.2 |
| Castilleja nana | 252.5 | 18.5 | 32.4 | -5.8 |
| Castilleja pilosa | 474.1 | 29.0 | 30.6 | -6.6 |
| Cerastium beeringianum | 168.1 | 20.8 | 31.8 | -6.4 |
| Chaenactis douglasii var. douglasii | 481.8 | 42.9 | 30.9 | -5.5 |
| Chamaebatiaria millefolium | 556.2 | 26.3 | 30.9 | -5.9 |
| Chrysothamnus viscidiflorus subsp. viscidiflorus | 584.5 | 21.4 | 33.6 | -3.9 |
| Cryptantha cinerea var. abortiva | 476.2 | 12.8 | 35.8 | -1.4 |
| Cryptantha flavoculata | 596.1 | 18.0 | 33.0 | -5.5 |
| Cryptantha humilis | 337.0 | 19.5 | 31.9 | -6.4 |
| Cymopterus cinerarius | 281.6 | 16.3 | 32.3 | -6.5 |
| Cystopteris fragilis | 531.1 | 39.1 | 32.1 | -3.5 |
| Dieteria canescens var. canescens | 549.9 | 21.3 | 32.7 | -4.8 |
| Draba albertina | 331.2 | 20.2 | 32.3 | -5.3 |
| Draba breweri | 213.7 | 16.2 | 33.8 | -4.3 |
| Draba californica | 184.5 | 14.1 | 32.7 | -6.4 |
| Draba oligosperma | 187.0 | 15.6 | 32.7 | -6.8 |
| Draba subumbellata | 159.6 | 14.3 | 33.0 | -6.7 |
| Eleocharis quinqueflora | 477.7 | 30.2 | 32.0 | -4.8 |
| Elymus cinereus | 648.6 | 30.5 | 30.2 | -6.7 |
| Elymus elymoides var. californicus | 364.2 | 61.6 | 29.0 | -5.7 |
| Elymus elymoides var. elymoides | 430.9 | 79.3 | 29.6 | -3.3 |
| Elymus sierrae | 235.9 | 19.5 | 31.7 | -6.8 |
| Eremogone kingii var. glabrescens | 320.2 | 20.8 | 31.9 | -6.4 |
| Ericameria discoidea | 258.8 | 17.6 | 32.6 | -5.4 |
| Ericameria suffruticosa | 277.8 | 18.9 | 31.9 | -6.5 |
| Erigeron clokeyi var. pinzliae | 382.1 | 15.9 | 32.3 | -6.2 |
| Erigeron compositus | 225.6 | 26.2 | 32.6 | -4.9 |
| Erigeron pygmaeus | 239.3 | 15.4 | 33.7 | -5.3 |
| Erigeron tener | 322.6 | 39.3 | 31.4 | -6.1 |
| Erigeron vagus | 112.8 | 16.7 | 32.7 | -5.7 |
| Eriogonum gracilipes | 250.9 | 14.3 | 32.5 | -5.8 |
| Eriogonum ovalifolium var. nivale | 280.0 | 16.7 | 33.2 | -5.3 |
| Eriogonum umbellatum var. dichrocephalum | 375.8 | 18.8 | 31.2 | -6.0 |
| Erythranthe suksdorfii | 432.2 | 21.1 | 33.5 | -3.6 |
| Festuca brachyphylla subsp. breviculmis | 164.9 | 15.7 | 32.9 | -5.6 |
| Festuca saximontana | 247.7 | 14.0 | 34.8 | -3.3 |
| Gayophytum decipiens | 501.2 | 20.7 | 33.0 | -3.4 |
| Gayophytum diffusum | 527.8 | 24.0 | 33.3 | -3.0 |
| Gymnosteris parvula | 276.6 | 16.3 | 32.2 | -6.5 |
| Heuchera parvifolia | 297.5 | 16.2 | 32.0 | -6.3 |
| Holodiscus discolor | 571.9 | 102.1 | 29.9 | -0.2 |
| Hulsea algida | 186.3 | 15.7 | 33.5 | -4.8 |
| Hymenoxys cooperi var. canescens | 345.9 | 16.2 | 32.6 | -5.7 |
| Ipomopsis congesta subsp. montana | 314.2 | 22.1 | 31.1 | -6.8 |
| Ivesia lycopodioides var. scandularis | 164.1 | 15.1 | 32.8 | -6.0 |
| Koeleria macrantha | 697.9 | 44.2 | 33.4 | -0.4 |
| Leptosiphon nuttallii | 365.0 | 57.4 | 30.9 | -5.5 |
| Lewisia pygmaea | 242.9 | 18.1 | 33.0 | -5.2 |
| Linanthus pungens | 577.4 | 24.2 | 33.7 | -2.6 |
| Linum lewisii var. alpicola | 345.8 | 14.4 | 34.0 | -5.0 |
| Lupinus argenteus var. heteranthus | 563.5 | 28.3 | 30.3 | -7.1 |
| Luzula spicata | 231.9 | 16.1 | 33.1 | -5.6 |
| Minuartia rubella | 228.8 | 24.6 | 33.5 | -4.2 |
| Monardella odoratissima subsp. glauca | 412.8 | 37.7 | 30.6 | -6.0 |
| Muhlenbergia richardsonis | 461.0 | 21.7 | 33.0 | -4.1 |
| Oxyria digyna | 236.6 | 29.1 | 32.4 | -5.0 |
| Oxytropis borealis var. viscida | 200.0 | 12.3 | 34.3 | -6.9 |
| Oxytropis parryi | 212.5 | 13.5 | 33.2 | -6.6 |
| Packera multilobata | 829.3 | 23.6 | 34.1 | -3.3 |
| Packera werneriifolia | 191.5 | 17.2 | 32.4 | -6.3 |
| Pedicularis attollens | 347.2 | 29.1 | 30.7 | -6.5 |
| Penstemon heterodoxus var. heterodoxus | 280.5 | 18.2 | 32.8 | -5.5 |
| Penstemon speciosus | 522.9 | 33.7 | 31.1 | -5.2 |
| Phacelia hastata var. compacta | 405.3 | 26.2 | 30.8 | -6.6 |
| Phlox condensata | 288.1 | 16.9 | 32.7 | -6.1 |
| Phlox hoodii subsp. canescens | 475.0 | 24.5 | 29.9 | -7.7 |
| Phlox pulvinata | 212.6 | 16.3 | 32.5 | -6.7 |
| Physaria kingii | 497.6 | 18.2 | 32.8 | -5.7 |
| Pinus flexilis | 447.6 | 18.1 | 34.8 | -1.4 |
| Pinus longaeva | 384.6 | 15.4 | 32.6 | -4.9 |
| Poa abbreviata subsp. pattersonii | 133.3 | 14.2 | 33.0 | -6.3 |
| Poa cusickii subsp. cusickii | 409.6 | 28.2 | 30.9 | -5.9 |
| Poa cusickii subsp. epilis | 277.5 | 25.4 | 31.6 | -6.0 |
| Poa cusickii subsp. pallida | 369.7 | 14.6 | 33.0 | -6.5 |
| Poa glauca subsp. rupicola | 195.2 | 16.7 | 33.1 | -5.9 |
| Poa keckii | 195.4 | 15.1 | 33.6 | -5.3 |
| Poa lettermanii | 129.8 | 16.7 | 32.9 | -4.0 |
| Poa secunda subsp. secunda | 724.5 | 49.5 | 32.2 | -2.2 |
| Polemonium chartaceum | 64.5 | 13.4 | 33.4 | -6.8 |
| Potentilla jepsonii | 276.9 | 12.1 | 34.6 | -6.6 |
| Potentilla morefieldii | 156.2 | 13.4 | 33.2 | -6.4 |
| Potentilla pensylvanica | 297.6 | 14.4 | 32.7 | -6.0 |
| Potentilla pseudosericea | 204.4 | 15.1 | 33.5 | -5.5 |
| Pyrrocoma apargioides | 291.3 | 16.1 | 32.9 | -6.0 |
| Ranunculus eschscholtzii | 220.2 | 30.6 | 32.2 | -4.7 |
| Ribes cereum var. cereum | 514.2 | 28.4 | 32.3 | -3.3 |
| Rumex paucifolius | 334.0 | 18.6 | 32.6 | -5.5 |
| Selaginella watsonii | 365.6 | 14.3 | 35.2 | -1.8 |
| Senecio integerrimus var. exaltatus | 437.1 | 36.0 | 30.4 | -6.3 |
| Silene bernardina | 450.6 | 33.2 | 32.3 | -4.5 |
| Solidago multiradiata | 303.6 | 27.4 | 32.2 | -5.4 |
| Sphaeromeria cana | 292.1 | 15.8 | 34.0 | -4.0 |
| Stenotus acaulis | 394.9 | 23.4 | 31.4 | -6.7 |
| Stipa hymenoides | 1031.4 | 15.6 | 36.6 | -0.7 |
| Stipa occidentalis var. occidentalis | 397.4 | 50.0 | 30.9 | -4.5 |
| Stipa pinetorum | 429.2 | 15.3 | 33.6 | -5.2 |
| Townsendia leptotes | 165.7 | 13.3 | 33.3 | -6.6 |
| Trifolium andersonii subsp. beatleyae | 237.0 | 13.7 | 32.9 | -6.4 |
| Woodsia scopulina subsp. scopulina | 144.3 | 16.4 | 31.9 | -7.2 |
