## Appendix S2 for "Community turnover by composition and climatic affinity across scales in an alpine system"

##### Smithers et al.— bioRxiv 2019 – Appendix S2

Appendix S2: Model results for each of the climate metrics on each of the peaks and for all peaks combined.

| **Climate metric** | **Location** | ***P-*value** | **F-df** | **F-value** | **Adj. R^2^** |
| --- | --- | --- | --- | --- | --- |
| **CWD** | **all** | **<0.001** | **1,15** | **69** | **80%** |
| **CWD** | **WMT** | **<0.001** | **1,11** | **31** | **71%** |
| **CWD** | **BAR** | **<0.001** | **1,12** | **24** | **64%** |
| CWD | SME | 0.313 | 1,9 | 1 | 1% |
| **CWD** | **PGS** | **0.001** | **1,6** | **32** | **82%** |
| CWD | CPT | 0.75 | 1,6 | 0 | 0% |
| **Annual Precip** | **all** | **<0.001** | **1,10** | **30** | **63%** |
| **Annual Precip** | **WMT** | **0.001** | **1,11** | **19** | **60%** |
| Annual Precip | BAR | 0.01 | 1,12 | 8 | 37% |
| Annual Precip | SME | 0.4 | 1,9 | 1 | 0% |
| Annual Precip | PGS | 0.253 | 1,6 | 2 | 0% |
| Annual Precip | CPT | 0.26 | 1,6 | 2 | 0% |
| **July T_max_** | **all** | **<0.001** | **1,22** | **49** | **72%** |
| July T_max_ | WMT | 0.0501 | 1,11 | 5 | 24% |
| **July T_max_** | **BAR** | **<0.001** | **1,12** | **23** | **63%** |
| July T_max_ | SME | 0.13 | 1,9 | 3 | 15% |
| **July T_max_** | **PGS** | **0.002** | **1,6** | **25** | **77%** |
| July T_max_ | CPT | 0.8 | 1,6 | 0 | 0% |
| **January T_min_** | **all** | **<0.001** | **1,20** | **34** | **64%** |
| January T_min_ | WMT | 0.15 | 1,11 | 2 | 10% |
| **January T_min_** | **BAR** | **0.001** | **1,12** | **18** | **56%** |
| January T_min_ | SME | 0.203 | 1,9 | 2 | 0% |
| January T_min_ | PGS | 0.04 | 1,6 | 6 | 43% |
| January T_min_ | CPT | 0.391 | 1,6 | 1 | 0% |
