## Appendix S3 for "Community turnover by composition and climatic affinity across scales in an alpine system"

##### Smithers et al.— bioRxiv 2019 – Appendix S3

##### Appendix S3: The 1961-1990 average climate variables (A) Climatic Moisture Deficit (CMD), (B) Mean Annual Precipitation (MAP), (C) July maximum temperature, and (D) January minimum temperature. For raster resolution purposes, we have here used CMD as a metric for water stress. CMD is positively correlated with climatic water deficit (CWD), which is used in analysis. Shading on the central range-wide linear line represents ±SE. Climate data was extracted from ClimateWNA v 5.51 (Wang et al., 2012).

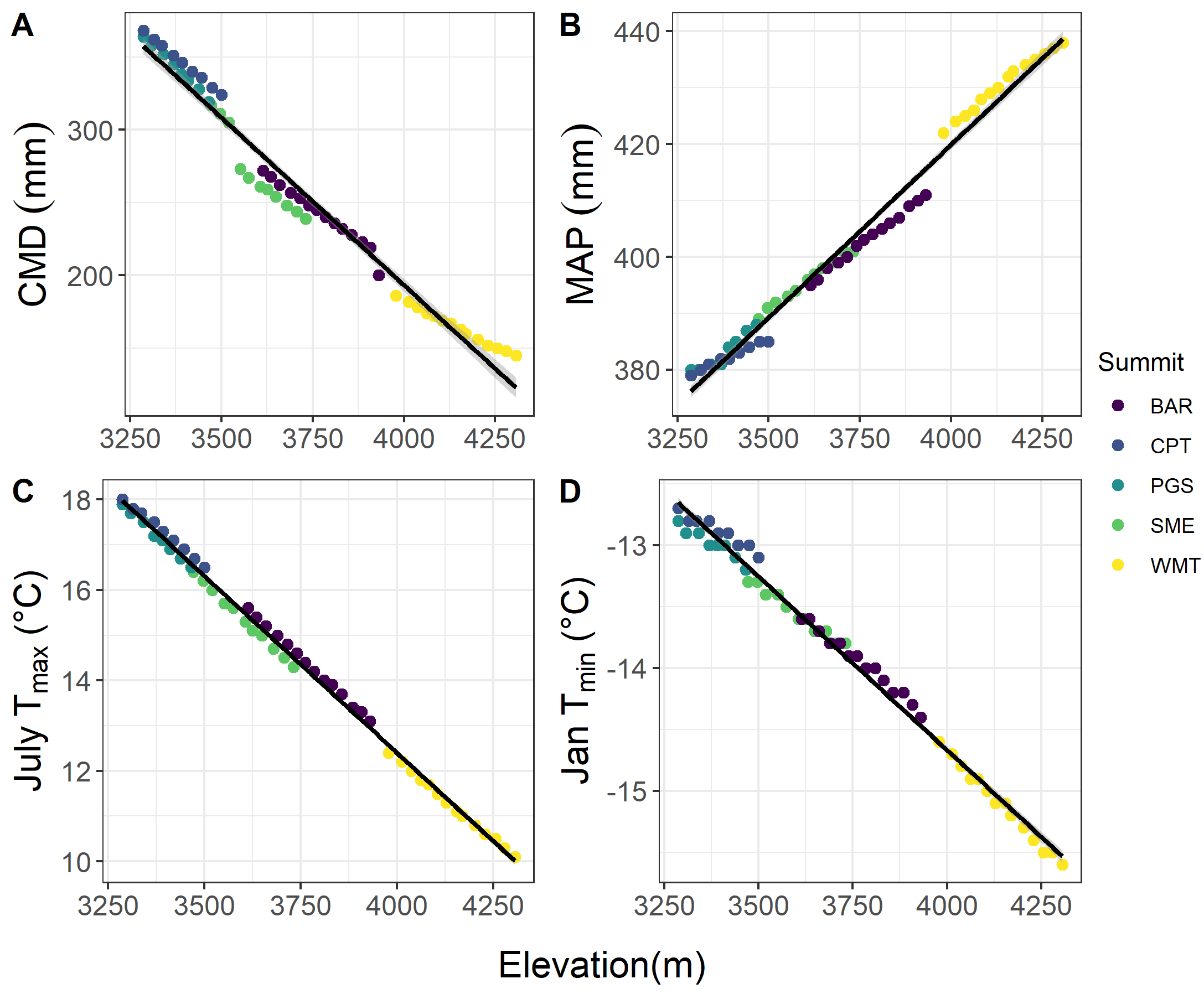
